# A Kinetic Map of the Influence of Biomimetic Lipid Membrane Models on Aβ_42_ Aggregation

**DOI:** 10.1101/2022.04.16.488542

**Authors:** Kevin N. Baumann, Michele Sanguanini, Oded Rimon, Greta Šneiderienė, Heather Greer, Dev Thacker, Matthias Schneider, Sara Linse, Tuomas P. J. Knowles, Michele Vendruscolo

**Affiliations:** University of Cambridge, Yusuf Hamied Department of Chemistry, Cambridge CB2 1EW, United Kingdom; University of Cambridge, Cavendish Laboratory, Cambridge CB3 0HE, United Kingdom; Lund University, Department of Biochemistry and Structural Biology, SE22100 Lund, Sweden

## Abstract

The aggregation of the amyloid β peptide (Aβ) is one of the major molecular hallmarks of Alzheimer’s disease. Although Aβ deposits have been mostly observed extracellularly, various studies have reported the presence of also intracellular Aβ assemblies. Because these intracellular Aβ aggregates might play a role in the onset and progression of Alzheimer’s disease, it is important to investigate their possible origins at different locations of the cell along the secretory pathway of the amyloid precursor protein (APP), from which Aβ is derived by proteolytic cleavage. Since lipid bilayers have been shown to promote the aggregation of Aβ, in this study we measure the effects of the lipid membrane composition on the in vitro aggregation kinetics of the 42-residue form of Aβ (Aβ_42_). By using small unilamellar vesicles modelling cellular membranes at different locations, including the inner and outer leaflets of the plasma membrane, late endosomes, the endoplasmic reticulum (ER), and the Golgi apparatus, we show that Aβ_42_ aggregation is inhibited by the ER and Golgi membranes. These results provide a preliminary map of the possible effects of the membrane composition in different cellular locations on Aβ aggregation, and suggest the presence of an evolutionary optimization of lipid composition to prevent the intracellular aggregation of Aβ.

## INTRODUCTION

The cytotoxic aggregation of Aβ is considered a major contributor to the onset and progression of Alzheimer’s disease (AD).^1–3^ While neurotoxic fibrillar deposits are found extracellularly, oligomeric species of Aβ are found intracellularly and in various cellular organelles, probably originating from a combination of local proteolytic processing, escape from the secretion pathway, and re-uptake of extracellular peptides.^4–7^ As Aβ is a product of the proteolysis of the membranebound amyloid precursor protein (APP), the influence of lipid membrane compositions on the aggregation of Aβ is of particular interest. Indeed, different membrane compositions have been shown to affect the proteolytic processing of APP and the kinetics of Aβ aggregation.^8–16^ Furthermore, cholesterol has been shown to increase the primary nucleation rate of Aβ in model cell membranes,^15^ an effect that is dependent on the composition of the lipid membranes themselves. It is also known that, while individual lipids can either accelerate or delay the aggregation of Aβ according to their physicochemical properties, mixtures of lipids can average out these effect and protect against aggregation aggregation.^16^ This phenomenon is of particular interest given that biological membranes exhibit a great repertoire of lipids and the membranes of organelles are constituted by individual and optimized lipid compositions.^17,18^ Therefore, because of the broad distribution of the individual effects lipid species composing a lipid membrane have on the aggregation of Aβ and other amyloids species,^16,19^ the most likely result is a neutral effect of that surface on Aβ. Yet, different compositions might result globally in either an enhancement or inhibition of the aggregation speed.

In this study we investigate the effects of lipid membrane compositions on Aβ_42_ aggregation using small unilamellar vesicles (SUVs) that mimic the composition of cellular components that have been implicated in Aβ processing and secretion: the outer and inner layer of the plasma membrane, late endosomes, the Golgi apparatus, and the ER. To obtain a physiologically relevant model for each lipid membrane, we selected the most prevalent lipid types as found in the literature^17,20^ (mol% > 5%, **Table 1**) and used complex mixtures of fatty acidic chains from commercially available purified brain lipid extracts. We analyzed the effects of the interaction of Aβ with the SUVs and the resulting aggregation by a combination of fluorescence assays and cryoelectron microscopy (cryo-EM).

**Table 1.** Lipid mixtures mimicking the lipid membrane compositions of different cellular compartments. Lipid compositions according to *: Lorent et al.^20^, **: Van Meer et al.^17^

| Cellular compartment | Lipid composition (mol%) |  |  |  |  |  |  |
| --- | --- | --- | --- | --- | --- | --- | --- |
|  | PC | PE | PI | PS | SM | BMP | Chol. |
| Outer leaflet of the plasma membrane* | 30 |  |  | 5 | 35 |  | 30 |
| Inner leaflet of the plasma membrane* | 22 | 40 |  | 26 |  |  | 12 |
| Late endosomes** | 40 | 15 |  |  | 7 |  | 27 |
| Golgi** | 43 | 17 | 9 |  | 17 |  | 14 |
| ER** | 52 | 26 | 9 |  |  |  | 13 |

Lipid types: PC (phosphatidylcholine), PE (phosphatidylethanolamine), PI (phosphatidylinositol), PS (phosphatidylserine), SM (sphingomyelin), BMP (bis(monoacylglycero)phosphate), chol (cholesterol).

## RESULTS AND DISCUSSION

The aggregation kinetics of Aβ_42_ in presence of different membrane-mimetic SUVs was monitored using the amyloid-sensitive dye thioflavin T (ThT) in a fluorescence assay (Methods). Our results show that lipid membrane compositions corresponding to the organelles involved early in the secretory pathway of APP inhibit the aggregation of the Aβ_42_ peptide (**Figure 1**), with aggregation half-time values for ER and Golgi that are up to double the ones of Aβ_42_ in the absence of SUVs. Conversely, the lipid membrane model of the outer leaflet of the plasma membrane accelerated the aggregation of the Aβ_42_ peptide by approximately 20% in comparison to native conditions. The lipid membrane models of the inner leaflet, on the other hand, exhibit a slightly inhibiting effect, and the membrane model of late endosomes denotes a tendency towards resilience or even enhancement of amyloid formation.

**Figure 1.**
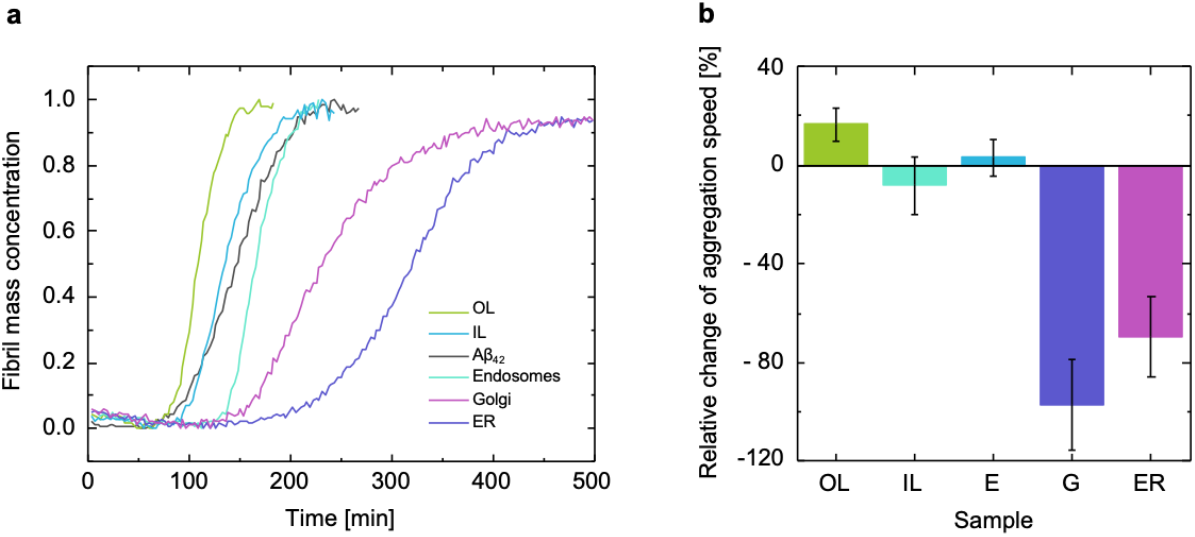
Aggregation kinetics of Aβ_42_ in presence of different lipid membrane models. **(a)** Set of representative traces of the Aβ_42_ aggregation kinetics in presence of the lipid membrane models visualized by a ThT fluorescence assay. The half time (t1/2) was calculated for each membrane composition and compared to that of Aβ_42_ in solution. **(b)** Using t1/2 as reference, the enhancement or inhibition of the aggregation kinetics imposed by the individual membrane compositions was measured. Error bars show the pooled standard deviation of at least 3 independent data sets per lipid membrane model. OL: outer leaflet of the plasma membrane, IL: inner leaflet of the plasma membrane, E: late endosomes, G: Golgi apparatus, ER: endoplasmic reticulum.

We next investigated whether Aβ_42_ aggregation modifies the morphology of the model lipid membranes. Using cryo-electron microscopy (cryo-EM), lipid membrane models of the outer leaflet, inner leaflet and endosomes presented a distinctly facetted surface, most likely due to the organization of the lipids into phase domains where specific lipid species dominate the local membrane composition (**Figure 2a-e**). More and longer fibrils were found in the samples containing the lipid membrane models of the outer leaflet and late endosomes, a possible effect of their effective enhancement of the aggregation kinetics. We found no morphological influence of the aggregation of the peptide on the SUVs, regardless of their lipid composition (**Figure 2f-j**). Furthermore, no or only weak association was found between the Aβ_42_ fibrils and the SUVs (**Figure 2 f-j**), in a way which resembles previous findings where TEM images of fibrils in presence of SUVs with complex lipid mixtures showed no morphological changes and only partial interactions between fibrils and SUVs.^10,16^

**Figure 2.**
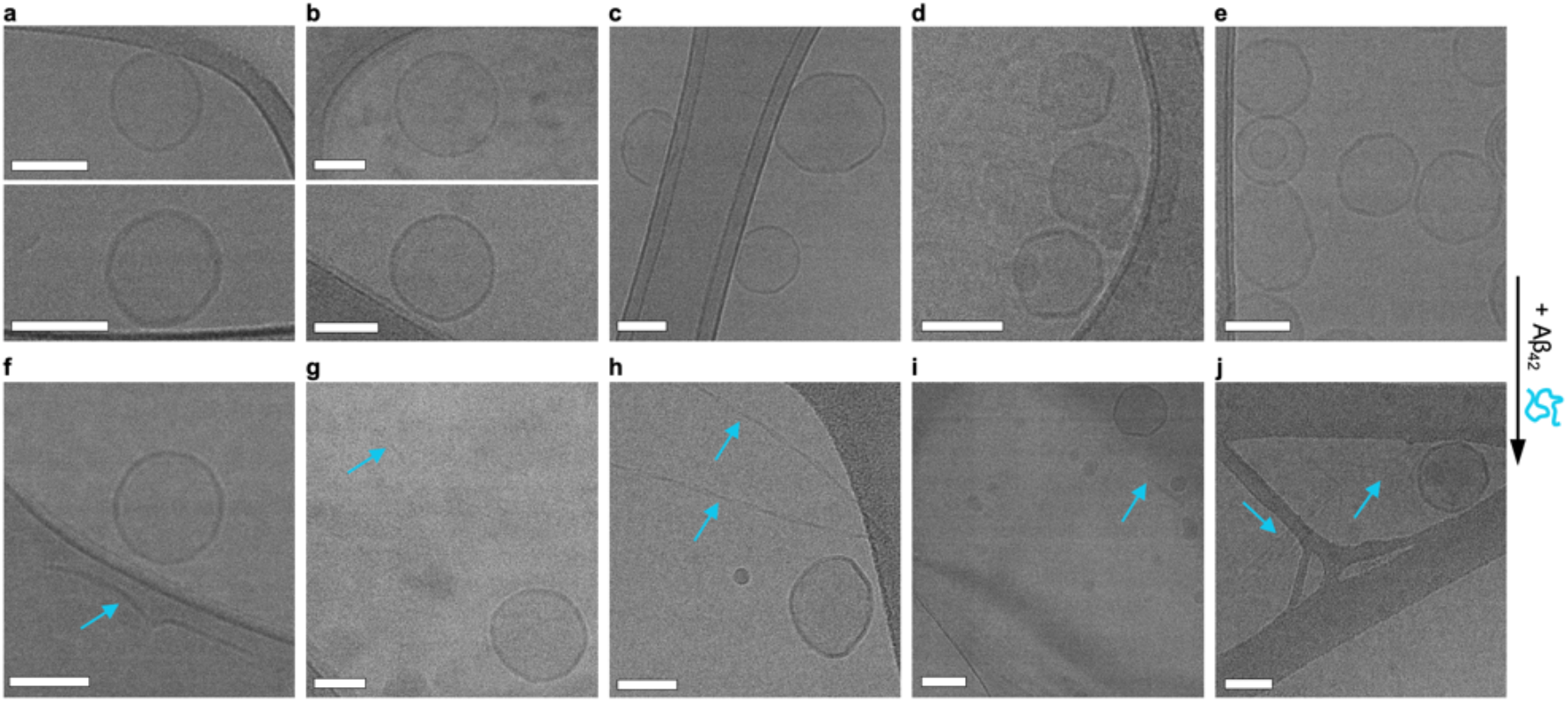
Cryo-electron microscopy reveals no morphological influence of Aβ_42_ aggregation on the SUVs. **(a-e)** Lipid membrane models of: Golgi (a), ER (b), endosomes (c), the inner (d) and outer leaflet (e) of the plasma membrane before the incubation with Aβ_42_ overnight. Endosomes, inner, and outer leaflet lipid membrane models show organization of the lipid membrane into facets. **(f-j)** After the addition of Aβ42, amyloid fibrils can be seen in all samples. Scale bars: 100 nm.

A possible explanation of the lack of strong interactions is that the SUVs facilitate the aggregation via a catalytic mechanism that involves the stabilization of the aggregation intermediates, but not of the reactants and the products of the aggregation reaction. According to this mechanism, the SUVs membranes might promote the aggregation of Aβ_42_ by transiently binding oligomeric species exposing extended hydrophobic surfaces, while not interacting permanently with the monomeric and the fibrillar forms, which expose fewer hydrophobic regions.^21–23^ By these means, oligomers bound to lipid membranes would dissociate less readily, thus having more time to undergo the conformational conversion step observed in the aggregation of Aβ_42_.^24^ In order to investigate this possibility, we asked whether lipid membranes induce a stable secondary structure of Aβ_42_, as one would expect in a scenario where lipid surfaces would induce peptide ordering.^25^ We monitored the evolution of circular dichroism (CD) spectra of Aβ_42_ in the presence of SUVs. Background-subtracted CD data show that Aβ_42_ does not change its secondary structure in the presence of varying concentrations of SUVs in the first hour (**Figure 3**). Within this time the peptide is expected to be present in a monomeric or early oligomeric state, according to the kinetic aggregation profiles. To further corroborate this possibility, we used microfluidic diffusional sizing to probe the binding of monomeric Aβ_42_ to the SUVs. Similarly to CD, no binding events were detected using this method (**Figure 4**). The experiments could not clarify conclusively whether or not small oligomers remain bound to the lipid membranes, as suggested by previous studies,^21,26^ and whether these bound oligomers are missing from an available pool of aggregation seeds, or if these further enhance the aggregation kinetics by initiating secondary pathways.^27^

**Figure 3.**
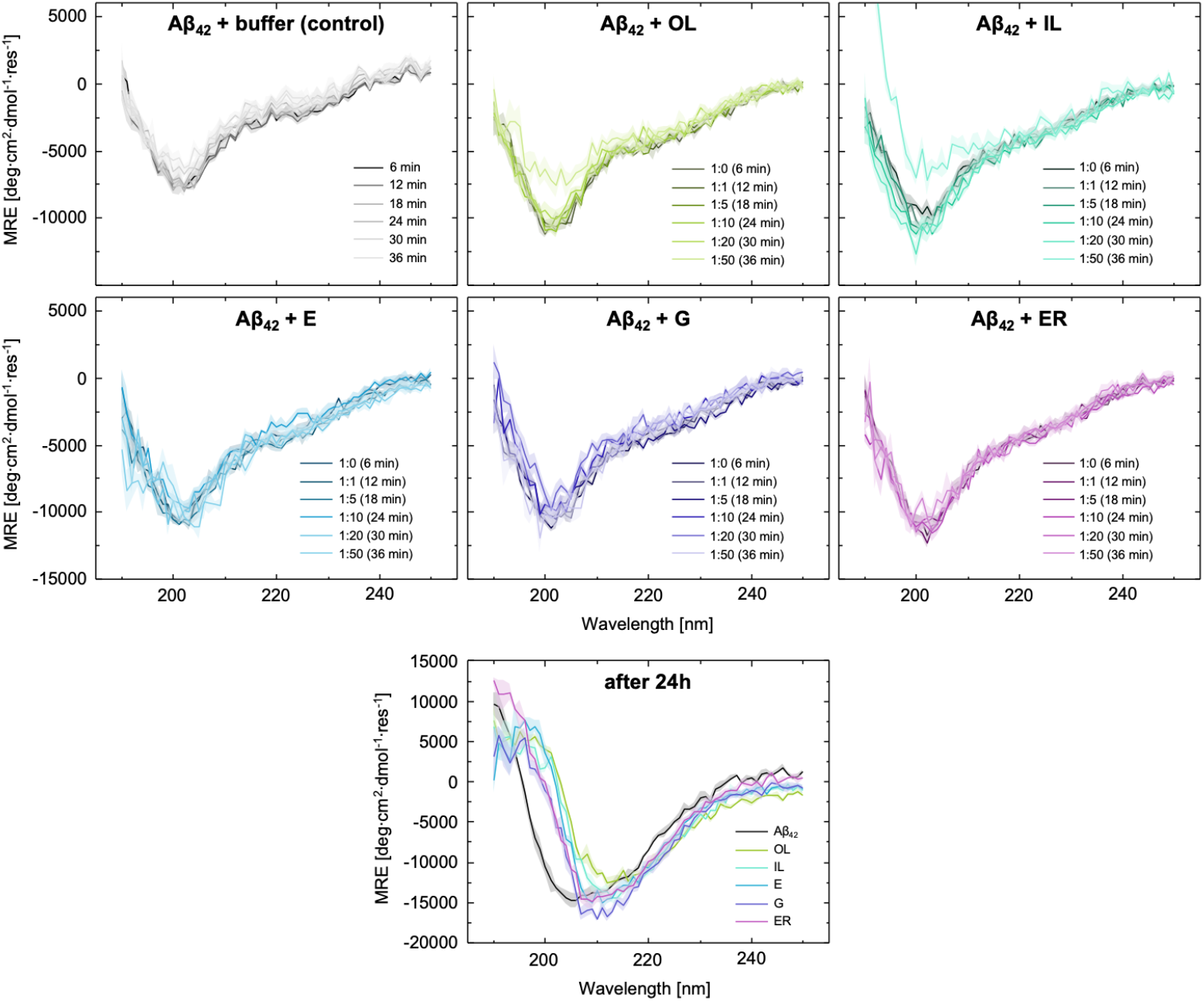
Aβ_42_ does not change its secondary structure in the presence of varying concentrations of SUVs. CD data of Aβ_42_ incubated with the model lipid membranes at increasing concentrations. After 24 hours (Aβ_42_ and SUVs ratio 1:50), the shift of the minimum by approximately 10 nm indicates β-sheet formation, enhanced by the lipid models.

**Figure 4.**
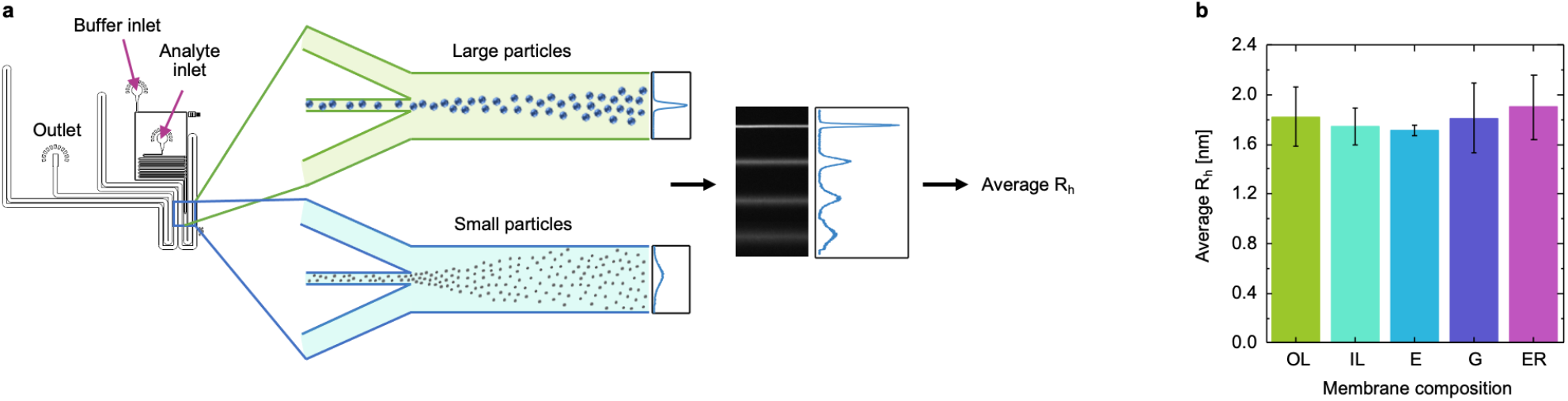
Microfluidic diffusional sizing measurements of the affinity of Aβ_42_ monomers to the lipid membrane models. Upon binding to a SUV, the average hydrodynamic radius (R_h_) of fluorescently-labelled Aβ_42_ is expected to increase by the radius of the SUV. However, no increase of the R_h_ of fluorescently-labelled monomeric Aβ_42_ upon mixing with the SUVs could be detected indicating that monomeric Aβ_42_ does not detectably interact with model lipid membranes. **(a)** Principle of the microfluidic diffusional sizing method: the higher diffusion rates of small particles lead to a broader distribution across the channel width at the detection region. **(b)** The average R_h_ of fluorescently-labelled Aβ_42_ monomers in mixture with the lipid membrane models was back-calculated based on these diffusion profiles. Error bars indicate the standard deviation (n =3).

In conclusion, our results indicate that the aggregation of Aβ_42_ is inhibited by model membranes mimicking the lipid composition of Golgi and ER membranes. The only membrane model that generated considerable enhancement of the aggregation kinetics was that of the outer leaflet of the plasma membrane. These observations support the theory of a possible evolutionary pressure towards the optimization of the membrane compositions of organelles in early stages of the APP secretory pathway to avoid the intracellular aggregation of Aβ_42_. More broadly, this study highlights the importance of understanding the intracellular toxicity of Aβ_42_ in its oligomeric form.^7,28^

## EXPERIMENTAL SECTION

### SUVs preparation

SUVs were prepared as previously described^16^ by extruding lipid solutions at 500 μM concentration suspended in a solution containing 20 mM NaHPO_4_ buffered with 0.2 mM EDTA (pH = 8.0) through 100 nm extrusion membranes (Avanti Lipids) after sonication at 40 °C for 30 min. All lipids, including those obtained from brain extracts, were purchased from Avanti and stored in chloroform. The SUVs compositions for the membrane models are gathered in **Table 1**.

### Aβ_42_ purification

The recombinant Aβ(M1-42) peptide (M DAEFRHDSGY EVHHQKLVFF AEDVGSNKGAIIGLMVGGVVIA), here referred to as Aβ_42_, was expressed in the *E. coli* BL21-Gold(DE3) strain (Stratagene, USA) and purified as described previously with slight modifications.^16,29^ Briefly, the transformed *E. coli* cells were sonicated, and the extracted inclusion bodies were dissolved in 8 M urea. The solution was then ion exchanged in batch mode on diethylaminoethyl cellulose resin and lyophilised. These lyophilised fractions were further purified using a Superdex 75 HR 26/60 column (GE Healthcare, USA), and the eluates were analysed using SDS-PAGE to confirm the presence of the desired protein product. The fractions containing the recombinant protein were pooled, aliquoted, frozen using liquid nitrogen, and lyophilised again to obtain the working stock.

### ThT assay

In order to prepare a solution of pure monomeric peptide, the lyophilized Aβ_42_ peptide was resuspended in 6 M guanidinium hydrochloride (GuCl) and then purified from excess salt and potential oligomeric species using gel filtration on a size exclusion column (Superdex 75 10/300 GL, GE Healthcare) at a flow rate of 0.5 mL/min and eluted in 20 mM sodium phosphate 0.2 mM EDTA buffer (at pH 8.0). The center of the peak was collected, and the peptide concentration was determined from the averaged concentration using the Lambert-Beer equation

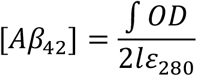

where *OD* is the optical density at 280 nm measured at the start and at the peak of the collection, *ε_280_* is the molar absorptivity coefficient at 280 nm (for Aβ_42_, ε_280_ = 1490 M·cm^-1^), and *l = 2* mm is the optical path length. The obtained peptide was diluted to the desired concentration with 20 mM sodium phosphate, 0.2 mM EDTA buffer (pH 8.0) and supplemented with 20 μM ThT and different molar-equivalents of SUVs. All samples were prepared in low binding test tubes (Eppendorf, Hamburg, Germany) on ice. Each sample was then pipetted into multiple wells of a 96-well half-area, low-binding, clear bottom, and PEG coating plate (Corning 3881, Corning, New York, NY, USA). Assays were initiated by placing the 96-well plate at 37 °C under quiescent conditions in a plate reader (Fluostar Omega or Fluostar Optima, BMG Labtech). The ThT fluorescence was measured through the bottom of the plate with a 440 nm excitation filter and a 480 nm emission filter.

### Cryo-electron microscopy

Cryo-EM grids were prepared by applying 3 μL of sample (at the native 1 nM concentration) on glow discharged holey gold grids (Quantifoil Cu 1.2/1.3 400 mesh). Excess sample was removed by blotting with filter paper for 4 sec prior to plunge-freezing in liquid ethane using a FEI Vitrobot Mark IV at 100% humidity and 4 °C. Data was collected on a FEI Tecnai F20 FEG microscope at 200 kV using a Falcon II direct electron detector. Images were collected at a dose rate of 20e^-^/pixel/second with a total dose of 20 e^-^/Å^2^. Magnification was set to 50,000x yielding a pixel size of 2.08 Å/pixel at the specimen level.

### Circular dichroism

Far-UV CD spectra were recorded between 190-250 nm using a Chirascan system (AppliedPhotopysics). A solution of 15 μM Aβ_42_ was transferred to a quartz cell with a 0.1 cm path length and incubated at 25 °C. After six minutes, a spectrum was recorded as θ (in mdeg). Following the initial measurement, a portion of buffer (control) or SUVs solution was added to achieve a 1:1 Aβ_42_:SUVs molar ratio, and six minutes after the first measurement the spectrum was recorded again. Every six minutes subsequently, a spectrum with increasing amounts of SUVs was recorded. Three spectra per time point were averaged, corrected by subtracting the buffer spectrum and normalised to Mean Residue Ellipticity (MRE; in deg · cm^2^ · dmol^-1^) using the sample concentration (in M) at that time point, and the number of residues in the protein

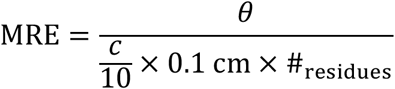

Finally, the spectra of all samples at 24 hours following the initial measurements (all at the final molar ratio of 1:50 Aβ_42_:SUVs) were recorded to reflect fully-aggregated controls.

### Aβ_42_ Y10C purification and labelling for microfluidic diffusional sizing

The Aβ_42_ Y10C mutant was purified as described above, except that 1 mM DTT was added to all buffers. Lyophilized fractions (~14 μM) of the peptide were dissolved in 50 μL deionized water. Alexa fluor 488 was added to the dissolved peptide in excess and kept overnight at 4 °C for labelling. The following morning, the mix was added in 1 mL of 6 M GuHCl, 20 mM sodium phosphate, 0.2 mM EDTA, pH 8.5 solution and subjected to gel filtration on a Superdex 75 10/300 column in 20 mM sodium phosphate buffer pH 8.0, with 0.2 mM EDTA. Absorption at wavelengths of 280 nm and 488 nm was monitored to follow the elution of the labelled peptide and to monitor any unlabeled peptide, if present. The aliquots collected from the SEC were then stored at −80 °C until further use.

### Microfluidic diffusional sizing

A master mold for the production of the polydimethylsiloxane (PDMS)-based microfluidic diffusional sizing devices generated using UV soft-lithography. Briefly, a negative photoresist (SU8-3050) was spin-coated on a silicon wafer to yield a 25–50 μm layer. The silicon wafer was then baked for 10 min at 95 °C on a hot plate. The wafer was then exposed to UV light through a photomask defining the channel geometries for 40 s. After exposure, the wafer was baked for 5 mins at 95 °C, followed by development in a propylene glycol methyl ether acetate (PGMEA) bath. The correct height of the features was measured with a profilometer.

To fabricate microfluidic devices, PDMS was mixed with carbon nanopowder (Sigma, USA) and a curing agent at a 10:1 mass ratio. The mixture was then centrifuged for 45 min at 5000 rpm, poured on the master and degassed under vacuum. Subsequently, the devices were baked for 1 h at 65 °C. Cured PDMS chips were then peeled off the master. A biopsy puncher was used to make channel inlets and outlets followed by device bonding to the glass slides. To this end, oxygen plasma treatment was used to activate the PDMS and glass surfaces.

The microfluidic diffusional sizing experiments were carried out as previously described.^30^ Before the measurements, the surface of microfluidic diffusional sizing devices was pre-treated with 0.01% Tween 20. 2.5 μM of Alexa 488-labelled Aβ_42_ Y10C mutant was mixed with 100 μM SUVs in a 20 mM NaHPO_4_, 0.2 mM EDTA (pH = 8.0) buffer and flown in a 25 μM diffusional sizing device at 50 μL/h flow rate. The devices were equilibrated for 5 mins before recording fluorescent traces across channels. To obtain the hydrodynamic radii, the images were analyzed with a custom-written Python script, utilizing the rate laws of diffusive mass transport under the laminar flow conditions.

### Dynamic light scattering

The SUVs were assessed regarding their monodispersity by dynamic light scattering (DLS) after production. The vesicles were measured at the native concentration using a ZetaSizer Nano (Malvern, UK) and Zen0040 disposable cuvettes. The measurements were performed at room temperature. The average hydrodynamic diameter of the SUVs was confirmed through DLS measurements. The values can be reviewed in the Supporting Information, **Figure S1**.

## Supporting information

Supplementary DLS results

## AUTHOR INFORMATION

### Author Contributions

The manuscript was written through contributions of all authors. All authors have given approval to the final version of the manuscript.

## ACKNOWLEDGMENTS

The authors would like to thank Thomas Löhr, Anne M. J. Jacobs, and Dillon Rinauro for insightful discussions.

## REFERENCES

(1) Knowles, T. P. J.; Vendruscolo, M.; Dobson, C. M. The Amyloid State and Its Association with Protein Misfolding Diseases. Nat. Rev. Mol. Cell Biol. 2014, 15 (6), 384–396. https://doi.org/10.1038/nrm3810.

(2) Hardy, J.; Selkoe, D. J. The Amyloid Hypothesis of Alzheimer’s Disease: Progress and Problems on the Road to Therapeutics. Science 2002, 297 (5580), 353–356. https://doi.org/10.1126/science.1072994.

(3) Hampel, H.; Hardy, J.; Blennow, K.; Chen, C.; Perry, G.; Kim, S. H.; Villemagne, V. L.; Aisen, P.; Vendruscolo, M.; Iwatsubo, T.; Masters, C. L.; Cho, M.; Lannfelt, L.; Cummings, J. L.; Vergallo, A. The Amyloid-β Pathway in Alzheimer’s Disease. Mol. Psychiatry 2021, 26 (10), 5481–5503. https://doi.org/10.1038/S41380-021-01249-0.

(4) Bu, G.; Cam, J.; Zerbinatti, C. LRP in Amyloid-Beta Production and Metabolism. Ann. N. Y. Acad. Sci. 2006, 1086 (1), 35–53. https://doi.org/10.1196/annals.1377.005.

(5) Wild-Bode, C.; Yamazaki, T.; Capell, A.; Leimer, U.; Steiner, H.; Ihara, Y.; Haass, C. Intracellular Generation and Accumulation of Amyloid β-Peptide Terminating at Amino Acid 42. J. Biol. Chem. 1997, 272 (26), 16085–16088. https://doi.org/10.1074/jbc.272.26.16085.

(6) Cook, D. G.; Forman, M. S.; Sung, J. C.; Leight, S.; Kolson, D. L.; Iwatsubo, T.; Lee, V. M. Y.; Doms, R. W. Alzheimer’s Aβ(1-42) Is Generated in the Endoplasmic Reticulum/Intermediate Compartment of NT2N Cells. Nat. Med. 1997, 3 (9), 1021–1023. https://doi.org/10.1038/nm0997-1021.

(7) LaFerla, F. M.; Green, K. N.; Oddo, S. Intracellular Amyloid-β in Alzheimer’s Disease. Nat. Rev. Neurosci. 2007, 8 (7), 499–509. https://doi.org/10.1038/nrn2168.

(8) Matsuzaki, K. How Do Membranes Initiate Alzheimers Disease? Formation of Toxic Amyloid Fibrils by the Amyloid β-Protein on Ganglioside Clusters. Acc. Chem. Res. 2014, 47 (8), 2397–2404. https://doi.org/10.1021/ar500127z.

(9) Hellstrand, E.; Sparr, E.; Linse, S. Retardation of Aβ Fibril Formation by Phospholipid Vesicles Depends on Membrane Phase Behavior. Biophys. J. 2010, 98 (10), 2206–2214. https://doi.org/10.1016/j.bpj.2010.01.063.

(10) Heo, C. E.; Park, C. R.; Kim, H. I. Effect of Packing Density of Lipid Vesicles on the Aβ42 Fibril Polymorphism. Chem. Phys. Lipids 2021, 236, 105073. https://doi.org/10.1016/j.chemphyslip.2021.105073.

(11) Niu, Z.; Zhang, Z.; Zhao, W.; Yang, J. Interactions between Amyloid β Peptide and Lipid Membranes. Biochim. Biophys. Acta - Biomembr. 2018, 1860 (9), 1663–1669. https://doi.org/10.1016/j.bbamem.2018.04.004.

(12) Sani, M. A.; Gehman, J. D.; Separovic, F. Lipid Matrix Plays a Role in Abeta Fibril Kinetics and Morphology. FEBS Lett. 2011, 585 (5), 749–754. https://doi.org/10.1016/j.febslet.2011.02.011.

(13) Choucair, A.; Chakrapani, M.; Chakravarthy, B.; Katsaras, J.; Johnston, L. J. Preferential Accumulation of Aβ(1-42) on Gel Phase Domains of Lipid Bilayers: An AFM and Fluorescence Study. Biochim. Biophys. Acta - Biomembr. 2007, 1768 (1), 146–154. https://doi.org/10.1016/j.bbamem.2006.09.005.

(14) Fabiani, C.; Antollini, S. S. Alzheimer’s Disease as a Membrane Disorder: Spatial Cross-Talk Among Beta-Amyloid Peptides, Nicotinic Acetylcholine Receptors and Lipid Rafts. Front. Cell. Neurosci. 2019, 13, 309. https://doi.org/10.3389/fncel.2019.00309.

(15) Habchi, J.; Chia, S.; Galvagnion, C.; Michaels, T. C. T.; Bellaiche, M. M. J.; Ruggeri, F. S.; Sanguanini, M.; Idini, I.; Kumita, J. R.; Sparr, E.; Linse, S.; Dobson, C. M.; Knowles, T. P. J.; Vendruscolo, M. Cholesterol Catalyses Aβ_42_ Aggregation through a Heterogeneous Nucleation Pathway in the Presence of Lipid Membranes. Nat. Chem. 2018, 10 (6), 673–683. https://doi.org/10.1038/s41557-018-0031-x.

(16) Sanguanini, M.; Baumann, K. N.; Preet, S.; Chia, S.; Habchi, J.; Knowles, T. P. J.; Vendruscolo, M. Complexity in Lipid Membrane Composition Induces Resilience to Aβ42 Aggregation. ACS Chem. Neurosci. 2020, 11 (9), 1347–1352. https://doi.org/10.1021/acschemneuro.0c00101.

(17) Van Meer, G.; Voelker, D. R.; Feigenson, G. W. Membrane Lipids: Where They Are and How They Behave. Nat. Rev. Mol. Cell Biol. 2008, 9 (2), 112–124. https://doi.org/10.1038/nrm2330.

(18) Vendruscolo, M. Lipid Homeostasis and Its Links With Protein Misfolding Diseases. Front. Mol. Neurosci. 2022, 0, 46. https://doi.org/10.3389/FNMOL.2022.829291.

(19) Ryan, T. M.; Griffin, M. D. W.; Teoh, C. L.; Ooi, J.; Howlett, G. J. High-Affinity Amphipathic Modulators of Amyloid Fibril Nucleation and Elongation. J. Mol. Biol. 2011, 406 (3), 416–429. https://doi.org/10.1016/j.jmb.2010.12.023.

(20) Lorent, J. H.; Levental, K. R.; Ganesan, L.; Rivera-Longsworth, G.; Sezgin, E.; Doktorova, M.; Lyman, E.; Levental, I. Plasma Membranes Are Asymmetric in Lipid Unsaturation, Packing and Protein Shape. Nat. Chem. Biol. 2020, 16 (6), 644–652. https://doi.org/10.1038/s41589-020-0529-6.

(21) Bode, D. C.; Freeley, M.; Nield, J.; Palma, M.; Viles, J. H. Amyloid-β Oligomers Have a Profound Detergent-like Effect on Lipid Membrane Bilayers, Imaged by Atomic Force and Electron Microscopy. J. Biol. Chem. 2019, 294 (19), 7566–7572. https://doi.org/10.1074/jbc.AC118.007195.

(22) Amaro, M.; Šachl, R.; Aydogan, G.; Mikhalyov, I. I.; Vácha, R.; Hof, M. GM1Ganglioside Inhibits β-Amyloid Oligomerization Induced by Sphingomyelin. Angew. Chemie - Int. Ed. 2016, 55 (32), 9411–9415. https://doi.org/10.1002/anie.201603178.

(23) Itoh, S. G.; Yagi-Utsumi, M.; Kato, K.; Okumura, H. Effects of a Hydrophilic/Hydrophobic Interface on Amyloid-β Peptides Studied by Molecular Dynamics Simulations and NMR Experiments. J. Phys. Chem. B 2019, 123 (1), 160–169. https://doi.org/10.1021/acs.jpcb.8b11609.

(24) Michaels, T. C. T.; Šarić, A.; Curk, S.; Bernfur, K.; Arosio, P.; Meisl, G.; Dear, A. J.; Cohen, S. I. A.; Dobson, C. M.; Vendruscolo, M.; Linse, S.; Knowles, T. P. J. Dynamics of Oligomer Populations Formed during the Aggregation of Alzheimer’s Aβ42 Peptide. Nat. Chem. 2020, 12 (5), 445–451. https://doi.org/10.1038/s41557-020-0452-1.

(25) Chi, E. Y.; Ege, C.; Winans, A.; Majewski, J.; Wu, G.; Kjaer, K.; Lee, K. Y. C. Lipid Membrane Templates the Ordering and Induces the Fibrillogenesis of Alzheimer’s Disease Amyloid-β Peptide. Proteins Struct. Funct. Genet. 2008, 72 (1), 1–24. https://doi.org/10.1002/prot.21887.

(26) Rushworth, J. V; Hooper, N. M. Lipid Rafts: Linking Alzheimer’s Amyloid-β Production, Aggregation, and Toxicity at Neuronal Membranes. Int. J. Alzheimers. Dis. 2011, 2011, 603052. https://doi.org/10.4061/2011/603052.

(27) Cohen, S. I. A.; Linse, S.; Luheshi, L. M.; Hellstrand, E.; White, D. A.; Rajah, L.; Otzen, D. E.; Vendruscolo, M.; Dobson, C. M.; Knowles, T. P. J. Proliferation of Amyloid-B42 Aggregates Occurs through a Secondary Nucleation Mechanism. Proc. Natl. Acad. Sci. U. S. A. 2013, 110 (24), 9758–9763. https://doi.org/10.1073/pnas.1218402110.

(28) Meli, G.; Lecci, A.; Manca, A.; Krako, N.; Albertini, V.; Benussi, L.; Ghidoni, R.; Cattaneo, A. Conformational Targeting of Intracellular A’ 2 Oligomers Demonstrates Their Pathological Oligomerization inside the Endoplasmic Reticulum. Nat. Commun. 2014, 5 (1), 1–17. https://doi.org/10.1038/ncomms4867.

(29) Walsh, D. M.; Thulin, E.; Minogue, A. M.; Gustavsson, N.; Pang, E.; Teplow, D. B.; Linse, S. A Facile Method for Expression and Purification of the Alzheimer’s Disease-Associated Amyloid β-Peptide. FEBS J. 2009, 276 (5), 1266–1281. https://doi.org/10.1111/j.1742-4658.2008.06862.x.

(30) Gang, H.; Galvagnion, C.; Meisl, G.; Müller, T.; Pfammatter, M.; Buell, A. K.; Levin, A.; Dobson, C. M.; Mu, B.; Knowles, T. P. J. Microfluidic Diffusion Platform for Characterizing the Sizes of Lipid Vesicles and the Thermodynamics of Protein-Lipid Interactions. Anal. Chem. 2018, 90 (5), 3284–3290. https://doi.org/10.1021/acs.analchem.7b04820.

