## Supplementary DLS results for "A Kinetic Map of the Influence of Biomimetic Lipid Membrane Models on Aβ_42_ Aggregation"

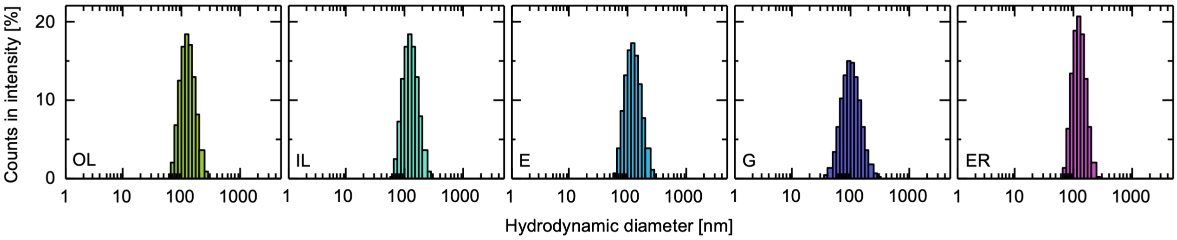


**Figure S1. Hydrodynamic diameters of the lipid membrane measured by dynamic light scattering**. OL: outer leaflet of the plasma membrane (130.7 nm ± 2.3 nm), IL: inner leaflet of the plasma membrane (129.6 nm ± 3.3 nm), E: late endosomes (127.8 nm ± 0.6 nm), G: Golgi apparatus (106.2 nm ± 6.9 nm), ER: endoplasmic reticulum (127.0 nm ± 2.9 nm). 3 independent SUVs preparations were averaged.
